# The Mystery of a Marine Monster: Morphological and Performance Modifications in the World’s only Marine Lizard, the Galápagos Marine Iguana

**DOI:** 10.1101/2020.05.16.099184

**Authors:** Kate A. Berry, Juan Pablo Muñoz-Pérez, Cristina P. Vintimilla-Palacios, Christofer J. Clemente

## Abstract

Reptiles have repeatedly invaded and thrived in aquatic environments throughout history, however fewer than 8% of the 6000 extant species are primarily aquatic. The Galápagos Marine Iguana (*Amblyrhynchus cristatus*), the world’s only marine lizard, may have had one of the most unique and challenging transitions to aquatic life. Curiously, previous studies have identified relatively few physiological adaptations in Marine Iguanas, however, little is known about the extent of morphological specialisation and performance trade-offs associated with the marine environment. By examining the morphology and locomotory performance of the Marine Iguana in comparison to their closely related mainland ancestors, the Black Spiny-tailed iguana (*Ctenosaura similis*) and Green Iguana (*Iguana iguana*), we found variation reflected specialisation to ecological niches. However, variation was more pronounced among subspecies of Marine Iguana, suggesting that little morphological or performance modification is required for iguanids to successfully invade aquatic environments, thus raising the question why there are so few extant aquatic reptilian lineages. We found that specialisation for the marine environment resulted in a trade-off in sprint speed in a terrestrial environment, similar to that seen in extant crocodilians. Reduced performance in a terrestrial environment likely poses little risk to large-bodied apex predators, whereas in iguanids, a performance trade-off would likely incur increased predation. As such, we suggest that this may explain why iguanids and other ancestral lineages have not undergone transitions to aquatic life. Additionally, we found that the magnitude of morphological and performance variation was more pronounced between subspecies of Marine Iguana than between iguanid species.

**Summary Statement:** The Marine Iguana has undergone a unique evolutionary transition to aquatic behaviour, we explore the extent of morphological and performance specialisation required and why there are so few extant marine reptiles.

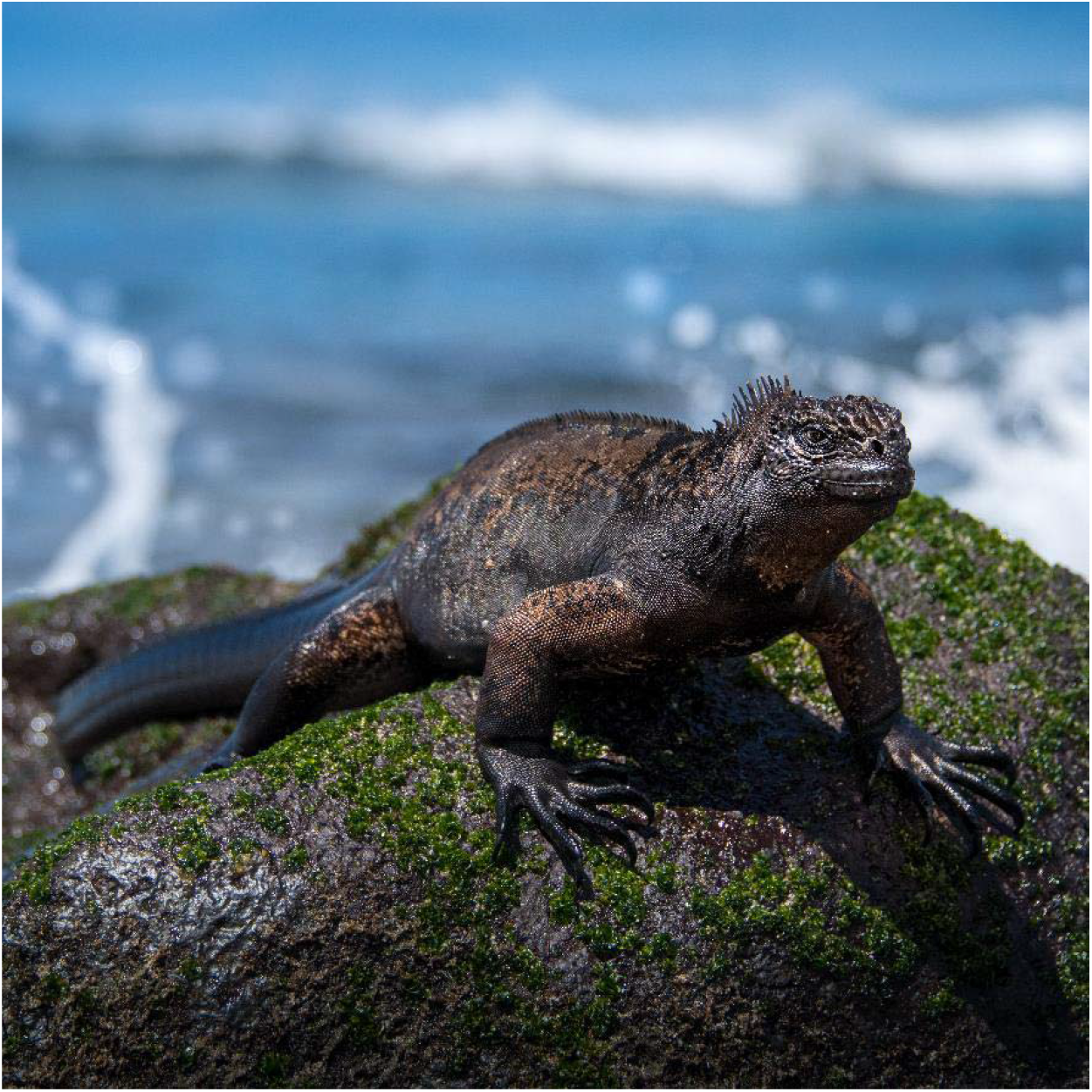

## Background

The extent to which an animal is adapted to the challenges of a specific habitat has long been a subject of evolutionary studies. The link between morphology, performance and fitness was first characterised in a framework described by Arnold (Arnold, 1983). This framework assumes that variation in morphology and physiology determines an individual’s fitness in a specific habitat only to the extent that it influences intermediary traits, such as locomotory performance(Arnold, 1983, Garland Jr and Losos, 1994). Under this view, natural selection acts most directly on locomotion, and therefore, locomotion is recognised as the foundation of all behaviour undertaken in an ecological niche (Mcelroy et al., 2007). The congruence between morphology, performance and habitat use is considered one of the most illustrative outcomes of adaptive evolution (Vanhooydonck and Van Damme, 2003). However, adaptations which promote performance in a single habitat can often reduce performance in other environments (Vanhooydonck and Van Damme, 2003). As such, trade-offs in performance are expected in animals that represent a transitional stage between terrestrial and aquatic life.

Reptiles are recognised as being one of the most evolutionary and ecologically remarkable components of global biodiversity. Reptiles not only have successfully invaded most areas of the world, they have also played a primary role in the origin and radiations of amniote vertebrates (Pincheira-Donoso et al., 2013). Consequently, reptiles are extensively used as model organisms in ecomorphological studies (Gomes et al., 2016). Although primarily terrestrial, reptiles have repeatedly invaded aquatic environments throughout history (Seymour, 1982). The Mesozoic era gave rise to at least a dozen diverse groups of marine reptiles, some of which were major oceanic predators for over 180 million years (Motani, 2009). Yet today fewer than 8% of the 6000 reptilian species are primarily aquatic (Bauer and Jackman, 2007). Previous studies have identified that the reptilian physiology allows for a simple transition into aquatic environments (Seymour, 1982, Motani, 2009), thus raising the question as to why so few lineages of aquatic reptiles exist today.

The Galápagos Marine Iguana (*Amblyrhynchus cristatus*) has arguably undergone one of the most significant evolutionary transitions among extant reptiles, being the world’s only marine lizard (Dawson et al., 1977). Marine iguanas must overcome numerous challenges associated with feeding exclusively on subtidal marine products, such as swimming through surf zones, overcoming buoyancy, tolerance of anaerobic conditions, the thermal constraints of diving in cold water and the exertion associated with underwater grazing (Seymour, 1982, Dawson et al., 1977). Surprisingly, Marine Iguana exhibit little departure from the physiological patterns characteristic of terrestrial iguanids (Dawson et al., 1977). For example, the species’ aerobic scope, thermal dependences, circulatory responses to diving, nasal salt secreting glands, reliance on anaerobiosis and restricted stamina all quantitatively resemble those of terrestrial iguanids (Dawson et al., 1977, Bartholomew and Lasiewski, 1965). Furthermore, previous studies have identified morphological modifications are often required for aquatic behaviour in reptiles, such as laterally compressed tails, short necks and webbed feet (Seymour, 1982, Motani, 2009). In fact, the tail and feet of Marine Iguana exhibited such little departure from the design of terrestrial iguanids that they did not warrant comment in the initial description of the species (Dawson et al., 1977). To date no studies have reappraised the morphological specialisations of Marine Iguana and quantified variation with terrestrial ancestors. Likewise, despite the central role performance plays in determining an individual’s fitness in a specific environment, little information exists on the locomotory ability of Marine Iguana in comparison to terrestrial ancestors.

In this study, we examine the morphology and locomotory performance of the Marine Iguana in comparison to their closely related mainland ancestors, the Black Spiny-tailed iguana (*Ctenosaura similis*) and the Green Iguana (*Iguana iguana*). This comparison explores the extent to which modification on the reptilian system is required for successful invasion of the marine environment, thus addressing how an iguanid evolved to become the world’s only marine lizard. Further, despite evidence that inter-specific studies require consideration of within-species variation, it is rarely considered in empirical works. As such, we investigate the magnitude of morphological and performance variation between newly identified subspecies of Marine Iguana as there is mounting evidence that the insular populations experience contrasting selection pressures.

## Methods

### Sampling

Morphometric and performance data was collected from 82 Marine Iguana across seven locations within the Galápagos archipelago in 2018. Sampling locations were selected based on Miralles et.al. (Miralles et al., 2017) taxonomic revision, resulting in data from seven populations representing six subspecies of Marine Iguana. Due to both geographic and genetic isolation described in McLeod et al. (MacLeod et al., 2015), we included two populations of *A.c.mertensi* in our subspecies analysis. A further 14 Black Spiny-tailed Iguana and 9 Green Iguana were collected from the Florida Keys in 2019. Data collection for this project was authorized by the Galápagos National Park Service (Permit #PC-86-18 - Kate Berry) and US Fish and Wildlife Service (FY 19-07). Ethics and animal handling protocols were approved by the University of the Sunshine Coast (#ANA18130), UNC-Chapel Hill and Universidad San Francisco de Quito (USFQ) Galápagos Science Center (GSC). Twenty morphological measurements, including limb, hip and head dimensions, were collected for each specimen (Figure 1).

**Figure 1:**
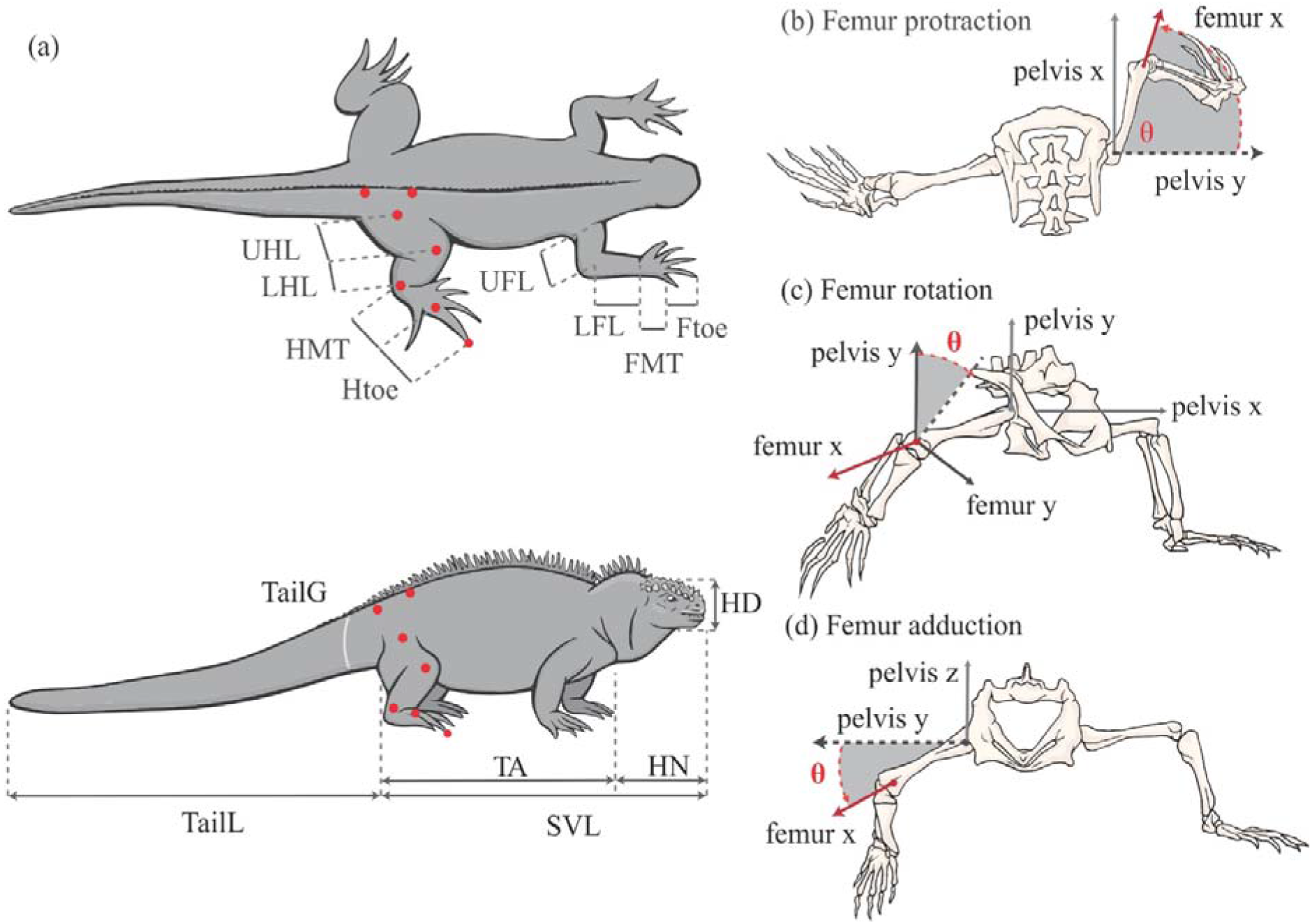
**a)** Morphological measurements and landmarks used in video digitisation. Also illustrated is the three movements of the femur relative to the pelvis including **b)** Femur protraction, **(c)** femur rotation and **(d)** femur retraction.

### Video Digitisation and Kinematics

Three-dimensional kinematics of the iguana’s stride, for each specimen was filmed running through a portable racetrack. Two Basler Ace USB3 Cameras (aca1920-150um, Basler, Ahrensburg, Germany), with either an 8mm or 16mm Tamron lens, were set on tripods perpendicular to the racetrack to capture both dorsal and lateral views of the specimens. Cameras were synchronised through custom recording software at 200 frames s^−1^, with a 1920 x 650 pixel resolution. To resolve spatial information, the racetrack space was calibrated using a two-point wand and DLT coefficients derived using easyWand software in the Matlab environment (Hedrick, 2008) (MATLAB 2017b, The MathWorks, Inc., Massachusetts, USA). Seven landmarks were painted on each iguana to mark the pelvis and hindlimb joints, to facilitate the video digitisation (Figure 1). Digitised landmarks were converted to three-dimensional coordinates using ‘DLTdv7’ digitising software within Matlab (Hedrick, 2008).

A total of 262 strides were digitised from videos of Marine Iguana (N=82, n=130), Black Spiny-tailed Iguana (N=14, n=69) and Green Iguana (N=9, n=44). Stride frequency (Hz) stride length (m), mean speed and maximal speed (m.s^−1^) was calculated for each stride. Further, twenty-five angular kinematic variables were calculated to described the movement of the hindlimb segments in relations to one another (Supp. Table S4). The maximum, minimum, midstance and the total angular change value was calculated for each kinematic variable.

### Analysis

Before analysis, mass, limb lengths and maximum speeds were log-transformed to normalise in R version 3.1.1 (Team, 2015). Analyses were performed firstly to identify patterns of variation among species, and secondly to identify patterns of variation among subspecies of Marine Iguana. Body size differences, speed variation, and speed modulation strategies were explored using lm.R and aov.R models in R where appropriate. The fastest two runs from each lizard were retained in speed analyses to reduce pseudo-replication. We calculated residuals for each length estimate from mass to remove any effects of body size and then explored shape variation using a linear discriminant analysis (LDA) with the lda.R function from the MASS package. Kinematic variables were explored using the lda.R function from MASS. Scores of the LDA were retained and used to estimate ancestral states using the fastANC.R function from the phytools package (Revell, 2012). The topology for phylogenetic trees was modified from (Miralles et al., 2017, Pyron et al., 2013).

## Results

### Body Size Variation in Iguanids

Body mass varied significantly between iguanid species (F_2,101_=10.39, P<0.0001; Supp. Figure S1). A Tukey Post Hoc test reported significant variation between Spiny-tailed Iguana and Marine Iguana (P<0.001). Marine Iguana had the greatest body mass (mean=2.18 ± 0.193, max=6.78kg), Green Iguana had intermediate (mean=0.582 ± 0.391, max=3.267kg), and Black Spiny-tailed Iguana the lowest masses (mean=0.312 ± 0.068, max=1.015kg). Patterns of SVL variation were consistent with those reported for body mass (F_2,101_= 10.04, P<0.001).

Body mass of male Marine Iguana also varied significantly among subspecies (F_6,32_=6.866, P <0.001; Supp. Figure S2). A Tukey Post Hoc test revealed a significant difference in body mass between the two populations of *A.c mertensi* (P<0.001), these being sampled from opposing coastlines of San Cristobal (Figure 2).

**Figure 2:**
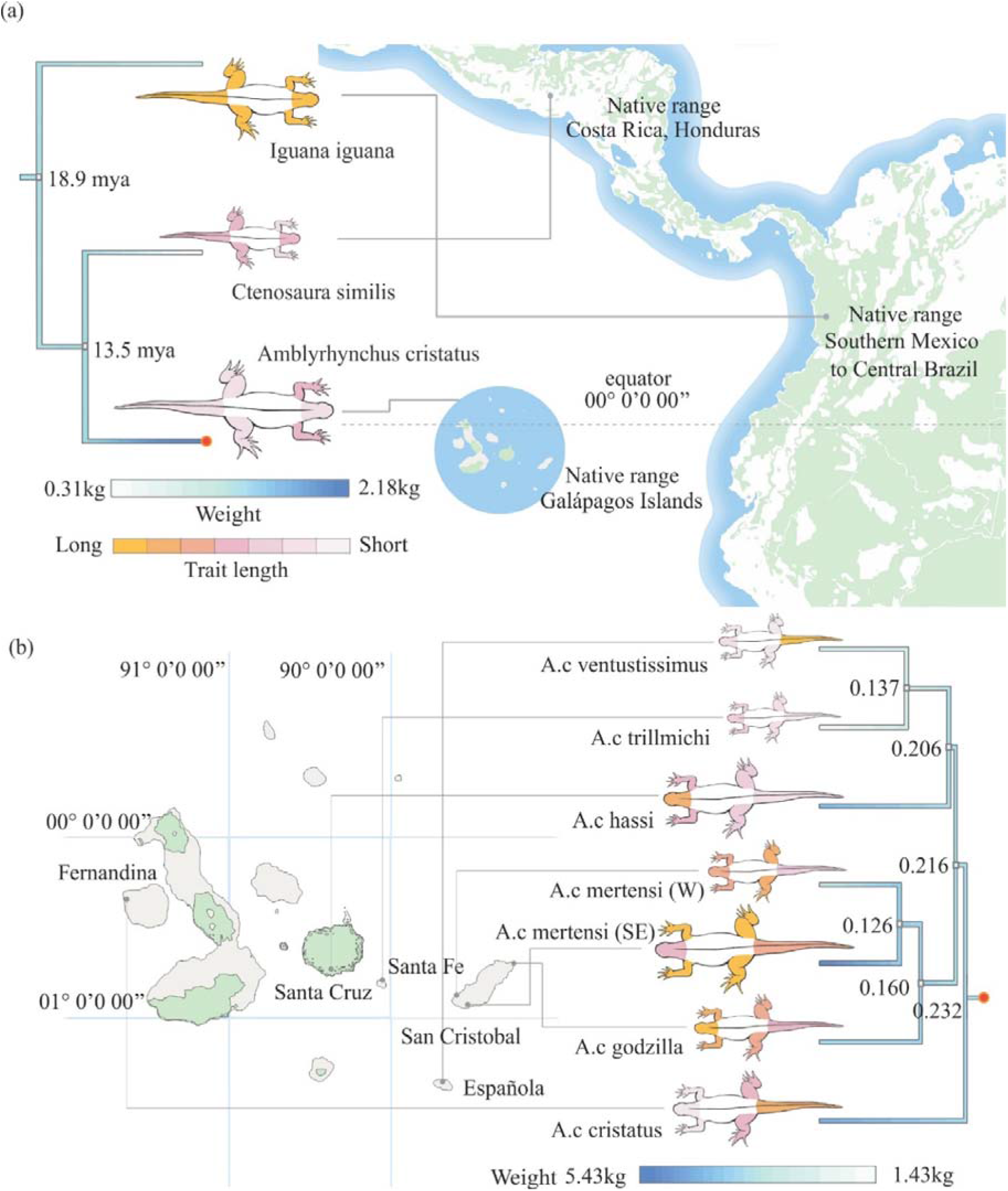
**(a)** Body shape and size variation between Marine Iguana, Green Iguana and Black Spiny-tailed Iguana. **(b)** Body size and shape variation in subspecies of Marine Iguana (juveniles excluded). Residual means of morphometric traits are expressed on the iguanid bodies. The ancestral state estimates for body mass are shown along phylogenetic trees. Morphometrics represented include head-neck length (HN), forelimb length (FL), hindlimb length (HL) and tail length (tailL). Iguanid bodies are scaled according to mean weight.

### Evolution of Body Shape in Iguanids

Variation in body shape, independent of size, was also visible among iguanids (Figure 2 & 3). An LDA significantly separated species, with the first function explaining 62.57% of variation (F_2,99_= 105.8, P<0.001). Along LD1 Marine Iguana and Spiny-tailed Iguana had similar scores, whereas Green Iguana were positive indicating long necks, tails, forelimbs, toes, and greatest hip depth (Supp. Table S1). The second discriminant explained 37.43% of shape variation, and separated Marine Iguana from mainland ancestors, by greater head depth, long forelimbs, shortest hindlimbs and widest hips.

**Figure 3:**
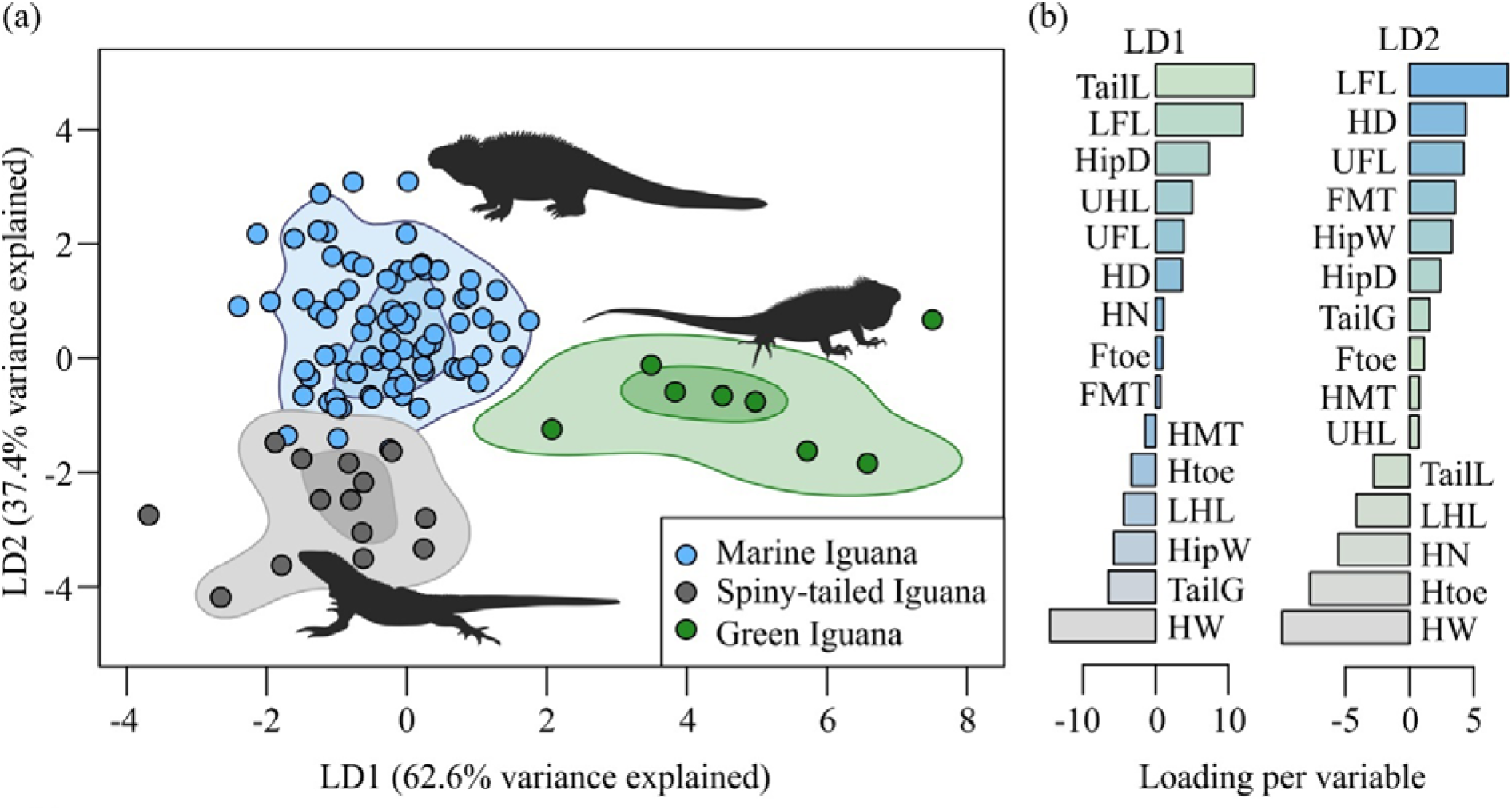
**(a)** A linear discriminant function illustrating shape variation between iguanids. Kernel density ellipses for each species illustrate 90% and 70% of the data distribution. **(b)** Graph of morphometric trait loadings from LD analysis.

Shape variation was also visible across subspecies of Marine Iguana (Figure 2). The first discriminant function explained 34.80% of variation between species (F_6,73_=12.83, P<0.001; Supp. Table S2) with a Tukey Post Hoc test indicating significant variation between *A.c mertensi* (western coast population) and all other subspecies (Figure 2).

### Evolution of Sprint Speed in Iguanids

There was no significant effect of mass on maximal speed in iguanids (F_1,90_=1.913, P=0.17), however there was a significant effect of species (F_1,90_=22.477, P<0.0001) and no significant interaction between the two (F_1,90_=1.032, P=0.36). Spiny-tailed Iguana had the greatest maximal speed recorded (Max 5.932 m.s^−1^, Mass = 0.44 kg), Green Iguana had intermediate speeds of 4.82 m.s^−1^ max, (Mass = 0.705kg) and Marine Iguana had the lowest maximal speed of 3.291 m.s^−1^ (Mass = 2.928kg; Supp. Figure S3).

Yet among Marine Iguana subspecies, there was a significant effect of mass on maximal speeds (F_6,61_=6.373, P=0.042), but no significant effect of subspecies (F_6,61_=2.090, P=0.067) with a significant interaction between the two (F_6,61_=2.325, P=0.043; Supp. Figure S4). The only significant difference in maximal speed reported in a Tukey Post Hoc test was between *A.c cristatus* and *A.c hassi* (P<0.001).

### Speed Modulation Strategies in Iguanids

Stride length significantly influenced speed in iguanids (F_1,228_=45.32, P<0.0001), but also varied among species (F_2,228_=47.56, P<0.0001) with a significant interaction between the two (F_2,228_=11.34, P<0.0001). A Tukey Post Hoc test revealed the Marine Iguana had the greatest mean stride length (0.508m), with no difference between the Green Iguana (0.391m) and Spiny-tailed Iguana, (0.409m; Figure 4). Similarly, stride length significantly affected speed among Marine Iguana subspecies (F_1,114_=34.242, P<0.001), with a strong effect of subspecies (F_6,114_=45.32, P=0.005) and no significant interaction of the two (F_1,114_=1.911, P=0.08). A Tukey Post Hoc test identified that the greatest variation in stride length was between *A.c trillmichi* and the south east coast population of *A.c mertensi*.

**Figure 4:**
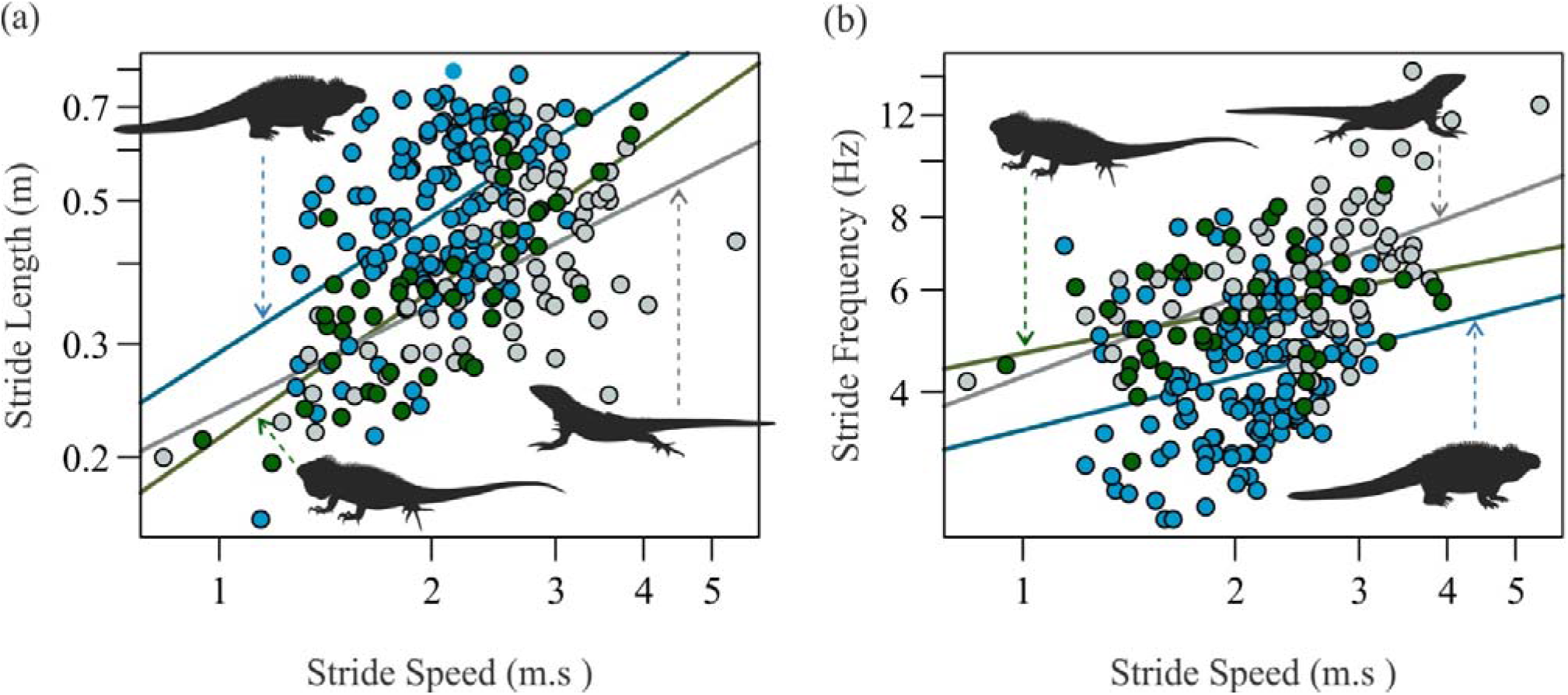
**(a)** Linear regression illustrating the relationship between log10 stride frequency and log10 stride speed for the three species of iguanids. **(b)** Linear regression illustrating the relationship between and log10 stride and log10 stride speed length for iguanids.

Both Stride frequency and species significantly influenced stride speed in iguanids (Freq: F_1,228_=76.55, P<0.001; Species: F_2,228_=8.970, P<0.005), with a significant interaction between frequency and species (F_2,228_=8.970, P<0.0001). Spiny-tailed Iguana had the highest stride frequency (mean=6.23 Hz), followed by Green Iguana (mean=5.40 Hz) and Marine Iguana (mean=4.16 Hz). Further, across Marine Iguana subspecies there was a significant effect of stride frequency on stride speed (F_6,114_=6.766, P<0.0001), a moderate effect on stride speed (F_1,114_=10.009, P<0.005) and no significant interaction (F_1,114_=0.861, P=0.526). A Tukey Post Hoc test reported the *A.c trillmichi* (5.648 Hz) was significantly higher than all subspecies except from *A.c cristatus* (P<0.001).

Combining the linear regression parameters from stride length and stride frequency indicated that iguanids modulate speed primarily by increasing stride length, however, the extent of use varied (Supp. Table S3). The intercept and slope reported that Marine Iguana take longer strides compared to their mainland ancestors. Intraspecific investigation revealed that this trend isn’t consistent across all Marine Iguana subspecies, with *A.c mertensi, A.c godzilla* and *A.c hassi* all modulate speed through primarily with their stride frequency.

### Iguanid Stride Kinematics

Variation in iguanid stride kinematics was visible in an LDA, where the first discriminant explained 88.86% of variation (F_2,228_=195.8, P<0.0001) and separated Marine Iguana from mainland ancestors. The second discriminant explained 11.34% of variation (F_2,228_=25.05, P<0.0001), with overlap in kinematics being observed among iguanids (Figure 5a). Overall, Marine Iguana exhibited the greatest knee angles (minimum and maximum), metatarsal angles (at midstance) and depression of the femur (maximum).

**Figure 5:**
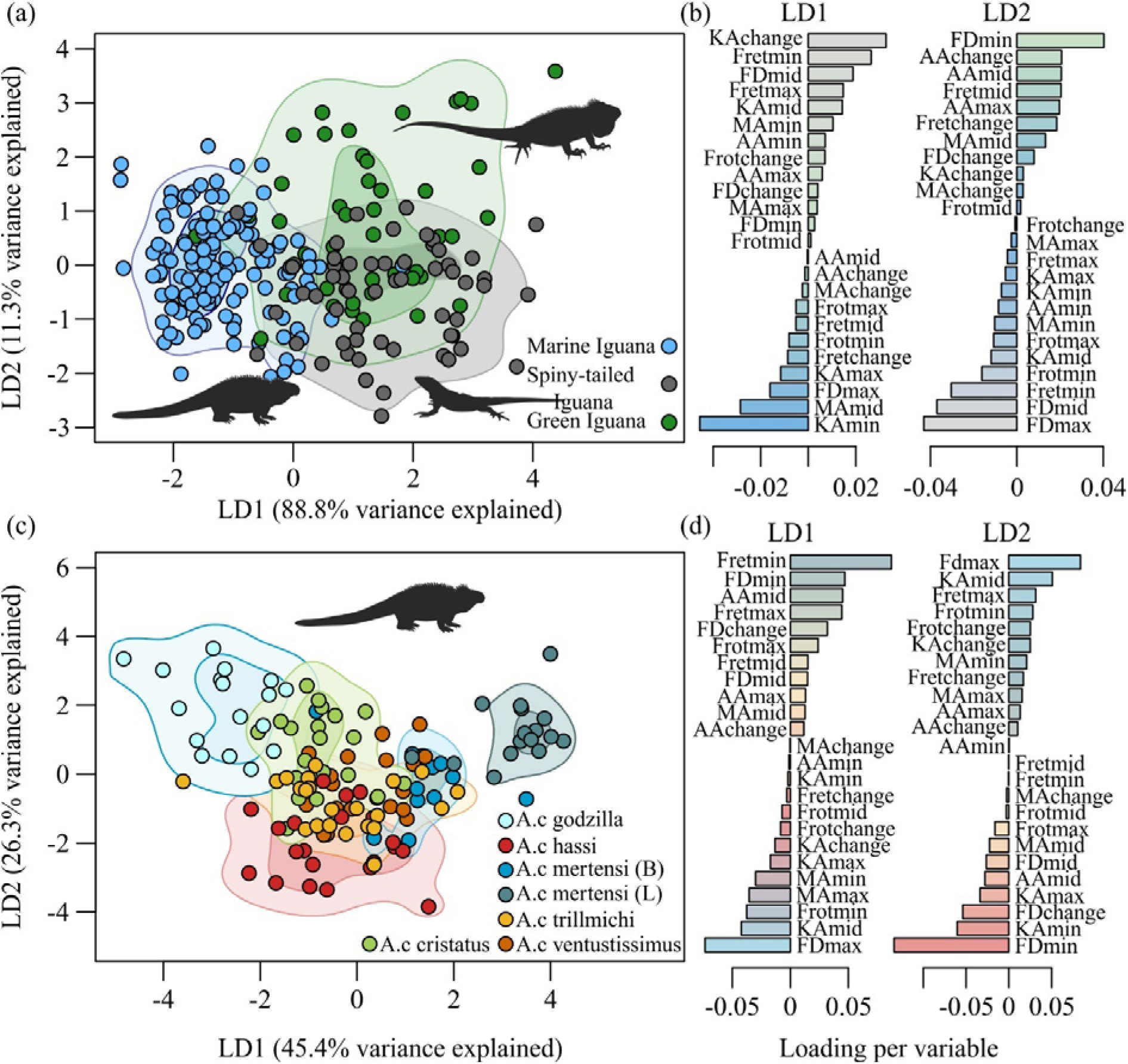
**(a)** LD function for variation in stride kinematics between iguanids, two of the fastest runs for each individual was retained (Marine Iguana N=65, Black Spiny-tailed Iguana N=35 & Green Iguana N=22). Kernel density ellipses for each species illustrate 90% and 70% of the data distribution. **(b)** Loadings for iguanid stride kinematics in (a). **(c)** LD function for variation in stride kinematics between subspecies of Marine Iguana. Kernel density ellipses as for (a). **(d)** Loadings for (c).

### Stride Kinematics in Marine Iguana Subspecies

Differences in stride kinematics between Marine Iguana subspecies were also evident, with the first discriminant explained 45.39% of the variation (F_6,121_=61.95, P<0.0001; Figure 5c). A Tukey Post Hoc test reported significant differences between *A.c godzilla* and *A.c hassi, A.c mertensi* (southeast and western populations), *A.c trillmichi* and *A.c ventustissimus*. The second discriminant explained 26.29% variation (F_6,121_=35.88, P<0.0001), with a Tukey Post Hoc test reporting significant variation between *A.c hassi* and *A.c cristatus*, as well as between *A.c godzilla* and *A.c trillmichi, A.c ventustissimus* and *A.c mertensi* (west coast population). Overall, the two populations of *A.c mertensi* varied from one another significantly (P<0.0001).

## Discussion

### Ecomorphological Associations in Iguanids

*Iguanidae* is one of the largest and most diverse families of lizards in the Western hemisphere, with species having radiated extensively into habitats ranging from rainforests to deserts, urban environments and rocky coastlines (Etheridge and de Queiroz, 1988). Studies suggest that specialisation for one habitat is generally assumed to occur at the cost of reduced fitness in another (Clemente et al., 2013, Vanhooydonck and Van Damme, 2003). As such, we hypothesised that the unique transition to a marine environment in iguanids would induce variation in morphology and performance trade-offs in Marine Iguana.

Body size is arguably the most fundamental design characteristic defining differences in closely related species and has been observed to change most readily on islands (Case and Schwaner, 1993, Petren and Case, 1997). Previous studies have shown islands harbour either gigantic or dwarf forms compared to mainland relatives, with size shifts often being a response to ecological pressures (Petren and Case, 1997). Our findings of greater body size in Marine Iguanas align with those of Petren and Case (Petren and Case, 1997), who reported that the island endemic chuckwallas, were up to five-fold larger in body mass compared to mainland species. Many theories exist that may explain the evolution of body size among iguanids. Shaw (Shaw, 1945) hypothesised that larger size was simply retained in genera that were first to diverge, whereas other studies suggest that recent evolution of small body size in mainland forms is consistent with a pattern of Holocene dwarfism (Case and Schwaner, 1993, Pregill, 1986). Invasive Green Iguana and Spiny-tailed Iguana were sampled from urban environments, therefore their smaller body sizes may be associated with anthropogenic effects. Yet, the shift to larger sizes in *Amblyrhynchus* is likely a response to island biogeography and depauperate environments (Case and Schwaner, 1993). Wikelski (Wikelski, 2005) proposed that evolution of size in Marine Iguana was a result of larger sizes being favoured because of the importance of thermal inertia on underwater foraging ability, as larger individuals are less sensitive to heat loss (Pincheira-Donoso et al., 2013).

Variation in shape also reflected specialisation for different environments. Marine Iguanas exhibited wide heads, blunt snouts, and short necks beneficial for underwater grazing and streamlined aquatic locomotion (Seymour, 1982, Motani, 2009). Long forelimbs are often associated with arborealism, as such we suggest that Marine Iguana have developed longer forelimbs to improve their manoeuvrability and stability when grazing underwater (Damme et al., 1998). In contrast, mainland iguanids had greater head neck lengths, longer tails, longer hindlimbs and toes than Marine Iguana, likely associated with varying degrees of arborealism. Previous studies have identified correlations between relatively shorter limbs and perch diameter in *Tropidurus, Draco* and Chameleons, hypothesised to be a trade-off between sprint speed and clinging ability (Kohlsdorf and Navas, 2012, Ord and Klomp, 2014, Hagey et al., 2017). As such, this trade-off likely explains differences in limb length between Spiny-tailed Iguana and Green Iguana. As Green Iguana are arboreal specialists it is relatively intuitive that they would have longer limbs, toes and tails as these traits benefit climbing ability and balance, whereas Spiny-tailed Iguana are only facultatively arboreal and thus do not require as notable specialisations. These morphological specialisations for aquatic behaviour may be associated with reduced performance in terrestrial environments (Clemente et al., 2013).

Sprint speed is used as a performance measure in ecomorphological studies as it can have a direct impact on fitness and is associated to limb morphology, body size, muscle fibre composition and the natural environment (Vanhooydonck et al., 2002). Our findings were consistent with existing literature, with the world record sprint speed for lizards having been attained by a Black Spiny-tailed Iguana (Garland Jr, 1984). It would be expected that a species from an open habitat, such as Spiny-tailed Iguana, would favour the evolution of traits that increase predator escape, such as greater sprint speed (Clemente et al., 2009). Whereas climbing species, such as Green Iguana, may be disadvantaged in terrestrial running as a result of trade-offs associated with their performance in vertical environments, such as clinging ability and sure-footedness (Clemente et al., 2013, Zaaf et al., 2001). Semi-aquatic reptiles may represent an intermediate phenotype between terrestrial and aquatic lifestyles, which would require biomechanical construction that enables locomotion on both land and in water (Seebacher et al., 2003). Therefore, we propose that navigation of rocky and wet substratum imposes conflicting demands on the locomotory ability of Marine Iguana, having likely occurred at the cost of reduced sprint speed in a terrestrial environment. This performance trade-off may be a contributing factor as to why so few extant reptilian lineages have invaded marine environments.

Sprint speed is a result of combining a particular stride length with a particular stride frequency, where an organism is able attain an identical speed solely by increasing either stride frequency or length (Clemente et al., 2013, Garland Jr and Losos, 1994). Iguanids were found to modulate speed primarily through changes in stride length, however the percentage of change varied subtly between species (Figure 4 & Supp. Table S3). Previous studies have found that ground-dwelling lizards achieve high speeds by increasing their stride length, whereas climbing species modulate speed through stride frequency (Clemente et al., 2009, Clemente et al., 2013). Therefore, the lack of difference in speed modulation strategies in iguanids, despite notable variation in sprint speed and habitat, suggests that predicted performance differences may not be as general as previously thought. Further, variation in the percentage of change in speed from stride length in iguanids may allow mainland ancestors to avoid lower speeds and overcome challenges of arborealism by adjusting stride frequency. Marine Iguana took significantly longer strides than mainland ancestors despite having relatively shorter hindlimbs. This association may be best explained by their stride kinematics and axial bending, which can serve to effectively increase hindlimb length.

Marine Iguana were found to increase knee, ankle and metatarsal angles, as well as femur rotation and retraction, all of which significantly contribute to an increase in effective limb length and thus an increase in stride length (Figure 5). An increase in effective limb length would increase the height of the body, which we hypothesise is to improve navigation of wet substratum. Reily and Elias (Reilly and Elias, 1998) hypothesised that semi-erect locomotion derived in alligators reflects the varying degrees of terrestrially. We observed striking similarities between the hip, femoral adduction and knee angles of *Alligator mississippiensis* and Marine Iguana (Blob and Biewener, 2001, Gatesy, 1997, Reilly and Elias, 1998). Therefore, our findings support that postural changes in reptiles may be associated with specialisation of aquatic environments (Reilly and Elias, 1998). Further, this effect may be disadvantageous to mainland iguanids, as lengthening limbs would hinder climbing ability by moving the body centre of mass away from the surface (Cartmill, 1985, Clemente et al., 2013, Zaaf et al., 2001). Unlike previous studies that have found changes in kinematics to be more pronounced than changes in morphology, we found overlap in iguanid joint angular changes (Clemente et al., 2013). This suggests that the increase in stride length in Marine Iguana may be more complex than being achieved solely through hindlimb joint angular changes. We hypothesise that lateral bending of the trunk, a common aspect of aquatic locomotion, may also increase their stride length (Ritter, 1992). Previous studies have identified that a travelling wave of lateral bending increases stride length and contributes to propulsive force of locomotion (Ritter, 1992). Therefore, we recommend investigation into the extent of lateral bending in Marine Iguana and hypothesise that it, along with a relative increase in hindlimb length through hindlimb joint changes, is likely the mechanism behind their increased stride length. Furthermore, we suggest that these biomechanical differences are associated with the demands of the marine environment, constraining terrestrial locomotory ability.

### Transitions to Aquatic Environments

Dawson et al. (Dawson et al., 1977) suggested that the physiological grade represented by the Iguanidae family was in a sense ‘pre-adaptive’ for aquatic habitats. Curiously, we found size and shape variation was more pronounced among subspecies of Marine Iguana than between iguanids, with some subspecies being up to four-fold larger in mass than others (Figure 2). Further, some subspecies were smaller than mainland ancestors, suggesting that increased size is not associated to the aquatic transition, and more likely associated with island biogeography and resource productivity (Wikelski, 2005, Wikelski and Trillmich, 1997). Historic studies also identified divergence in morphology between different populations of Marine Iguana (Chiari et al., 2016, Wikelski, 2005, Wikelski and Trillmich, 1997), although the ecological considerations of such variation requires further investigation. Interestingly, *A.c mertensi, A.c godzilla* and *A.c hassi* were found to modulate speed primarily with their stride frequency, thus differing to the primary strategy used by iguanids (Supp. Table S3). Further evidence for significant subspecies diversification was the extent of variation in their stride kinematics, differing on very localised scales, such as between populations of *A.c mertensi* from opposing coastlines (Figure 5c). These intraspecific findings demonstrate that the morphological and performance variation between iguanids would not inhibit the initial transition to aquatic life, thus raising the question why so few reptiles have exploited aquatic environments.

We found evidence that specialisation to the marine environment in Marine Iguana has led to a trade-off in their performance (sprint speed) in terrestrial environments. Similarly, extant crocodilians, despite having developed an array of movement patterns (high walk and gallop), have reduced locomotory performance in terrestrial environments (Seebacher et al., 2003, Webb and Gans, 1982). Reduced performance in a terrestrial environment likely poses little risk to large-bodied apex predators, whereas in iguanids, a performance trade-off would likely incur increased predation, as nearly all continents are occupied by mammalian predators (Dawson et al., 1977). Studies have suggested that the success of Marine Iguana is largely due to the lack of predation in Galápagos, where there are few consequences of being slow on land (Berger et al., 2007). Therefore, we suggest that other iguanids have not invaded aquatic environments due to the trade-off in their terrestrial performance, which may be detrimental to survival.

Further, the unique biogeography of the Galápagos may also explain why other lizards have not invaded marine environments. The equatorial position of the Galápagos archipelago, and minimal variation in seasons, is an important ecological consideration for heliothermic lizards (Seymour, 1982). Continuous upwelling of nutrient-rich, cold waters, containing soft-bodied macrophytic algae, is quite localised, with comparable upwelling regions found along the coasts of Nambia, Chile, California and Northwest Africa, all being outside the geographic range of Iguanids (Berger et al., 2007). Furthermore, lizard diversity is low in all of these regions and the presence of mammalian predators likely explains why other species have not invaded marine environments in a similar manner to that of the Marine Iguana.

### The Mysteries of Marine Iguana – Subspecies Diversity

Oceanic islands have long been recognised as natural laboratories for the study of evolution, as significant genetic and morphological variation often occurs as a consequence of the exploitation of free niches or multiple colonisation events (Van Valen, 1965, Losos et al., 1998). Studies have documented a range of body size within Marine Iguana, however, majority of findings have been based off samples collected over two decades ago (Wikelski, 2005, Wikelski and Thom, 2000, Wikelski and Trillmich, 1997, Chiari et al., 2016, Miralles et al., 2017). Surprisingly, we found that subspecies of Marine Iguana exhibited more variation in size, shape and performance than between iguanids, despite occupying similar niches across the Galápagos.

We found significant variation in body size between subspecies of Marine Iguana, with some subspecies being up to four-fold larger in mass than others. Further, some subspecies were smaller than mainland ancestors, suggesting that increased size is not associated to marine life and supports greater association with island biogeography and resource productivity (Wikelski, 2005).

Despite our interspecific findings suggesting that island biogeography cause shifts to larger body sizes in iguanids, patterns of size variation in Marine Iguana varied significantly from historical records documented by (Chiari et al., 2016). We found there has been a six-fold reduction in body size in Marine Iguana over the past two decades. In contrast, (Wikelski, 2005) found that size increased over the 19^th^ century. Over the past two decades, there have been two strong El Niño events (1997-1998 and 2015-2016), which caused periodic disappearance of the iguana’s preferred food source and thus population crashes and depressed growth rates (Laurie, 1990, Romero and Wikelski, 2002). Larger body sizes are strongly selected against during El Niños, whereas sexual selection drives an increase in size (Wikelski, 2005). During the 1997-1998 El Niño, individual Marine Iguana temporarily shrunk up to 20% of their body length, hypothesised to be an adaptive response to low food availability (Wikelski and Trillmich, 1997). We suggest that over time, climatic variation has had a more pronounced effect on body size than sexual selection. As the frequency and severity of El Niño events increase, we suspect that this selection will continue to favour a shift towards smaller sizes across the archipelago (Cai et al., 2014). Consequently, the shift in body size is complex, historically being driven by island biogeography, whereas now, climate is likely having a more pronounced effect on the patterns.

Historic studies also identified divergence in morphology between different populations of Marine Iguana (Chiari et al., 2016, Miralles et al., 2017). Similar to our findings in body size, morphology and locomotory ability was in some instances more varied among Marine Iguana subspecies than between iguanid species, with *A.c mertensi, A.c godzilla* and *A.c hassi* modulating speed primarily with stride frequency. However, modulating speed with stride frequency did not appear to be associated with the highest attainable sprint speeds in Marine Iguana. Subspecies found on San Cristobal also had notable differences in their stride kinematics, despite records of hybridisation and divergence times (MacLeod et al., 2015). Therefore, we suggest that variation in design correlates with performance differences in Marine Iguana, which is likely associated to localised, structural differences in habitat. This illustrates that the form-function relationship is complex and highlights the ecomorphological mysteries of Marine Iguana. Further investigation is recommended into microhabitat use by Marine Iguana to determine the mechanisms behind their unique variation.

## Acknowledgements

We are grateful to the Galápagos National Park authority, Florida Fish and Wildlife Conservation Commission and Florida Keys National Wildlife Refuges for permitting data collection for this project; to the UNC-USFQ Galápagos Science Centre for facilitating fieldwork and laboratory space; and to the volunteers and collaborators that assisted during data collection in Galápagos and Florida. Thank you to the National Geographic Society and the University of the Sunshine Coast for funding this research.

## Competing Interests

All authors hereby declare that they have no competing interests.

## Funding

This work was supported by a National Geographic Society grant awarded to K.A.B (ec-327r-18), an ARC discovery grant awarded to C.J.C (DP180100220) and by the University of the Sunshine Coast.

## Data Accessibility

Data will be submitted along the manuscript via Figshare. https://doi.org/10.6084/m9.figshare.12121449.v2

